# Prediction in Human Auditory Cortex

**DOI:** 10.1101/474718

**Authors:** KJ Forseth, G Hickok, Patrick Rollo, N Tandon

## Abstract

Spoken language is thought to be facilitated by an ensemble of predictive mechanisms, yet the neurobiology of prediction for both speech perception and production remains unknown. We used intracranial recordings (31 patients, 6580 electrodes) from depth probes implanted along the anteroposterior extent of the supratemporal plane during rhythm listening, speech perception, and speech production. This revealed a frequency-multiplexed encoding of sublexical features during entrainment and a traveling wave of high-frequency activity across Heschl’s gyrus. Critically, we isolated two predictive mechanisms in early auditory cortex with distinct anatomical and functional characteristics. The first mechanism, localized to bilateral Heschl’s gyrus and indexed by low-frequency phase, predicts the timing of acoustic events (“when”). The second mechanism, localized to planum temporale in the language-dominant hemisphere and indexed by gamma power, predicts the acoustic consequence of speech motor plans (“what”). This work grounds cognitive models of speech perception and production in human neurobiology, illuminating the fundamental acoustic infrastructure – both architecture and function – for spoken language.

## Introduction

Humans efficiently extract speech information from noisy acoustic signals and segment this into meaningful linguistic units. This complex and poorly understood process is fluidly accomplished for a wide range of voices, accents, and speaking rates^1,2^. Given the quasi-periodic nature of speech^3^, the computational load associated with its decoding can be reduced by utilizing temporal prediction. Anticipating the arrival of salient acoustic information enables optimal potentiation of neural networks^4–7^ and discretization of the continuous signal into linguistic elements^4,8,9^. Evidence for cortical entrainment – the synchronization of extrinsic pseudo-periodic stimuli and intrinsic neural activity – to speech^10–14^ have driven speculation that cortical oscillations may encode temporal prediction.

The production of speech is another human ability that relies on predictive mechanisms. There is now strong evidence that speech planning involves prediction of the sensory consequences of the action^15–20^. It remains unclear, however, which levels of auditory cortical processing are involved in this process and where such mechanisms are instantiated in the cortex.

The identification and analysis of predictive mechanisms for language function requires a methodology with high temporal resolution, fine spatial resolution, and direct access to neuronal populations in human early auditory cortex. We used large-scale intracranial recordings (31 patients, 6580 electrodes), focusing on depth electrodes placed in an innovative trajectory along the anteroposterior extent of the supratemporal plane. We investigated cortical entrainment and prediction during a novel amplitude-modulated white noise stimulus, as well as during natural language speech. These experiments yield crucial insights into the rapid, transient dynamics of prediction for “when” and “what” in Heschl’s gyrus and planum temporale.

## Results

### Entrainment to Low-Level Acoustic Stimuli

Recordings along the supratemporal plane revealed entrainment of early auditory cortex to rhythmic amplitude-modulated white noise (80% depth at 3 Hz for 3 seconds, then constant amplitude for 1 second; Figure 1A). Heschl’s gyrus and the transverse temporal sulcus (HG/TTS; Figure 1B) encoded stimulus features in gamma power (65-115 Hz, Figure 1C) and low-frequency phase (2-15 Hz, Figure 1D). Following a low-latency high-magnitude broadband response to stimulus onset, this region entrained to subsequent acoustic pulses. Phase space trajectories of gamma power (Figure 1G) and low-frequency phase (Figure 1H) revealed three clearly dissociable states corresponding to rest (pre-stimulus), stimulus onset, and entrainment (beginning with the second pulse). These results were robust across the patient cohort; at least one electrode recorded an acoustic entrainment response in all patients with a supratemporal depth probe in the language dominant hemisphere (n = 18). Patients with homologous electrodes in the language non-dominant hemisphere (n = 4) demonstrated equivalent entrainment.

**Figure 1:**
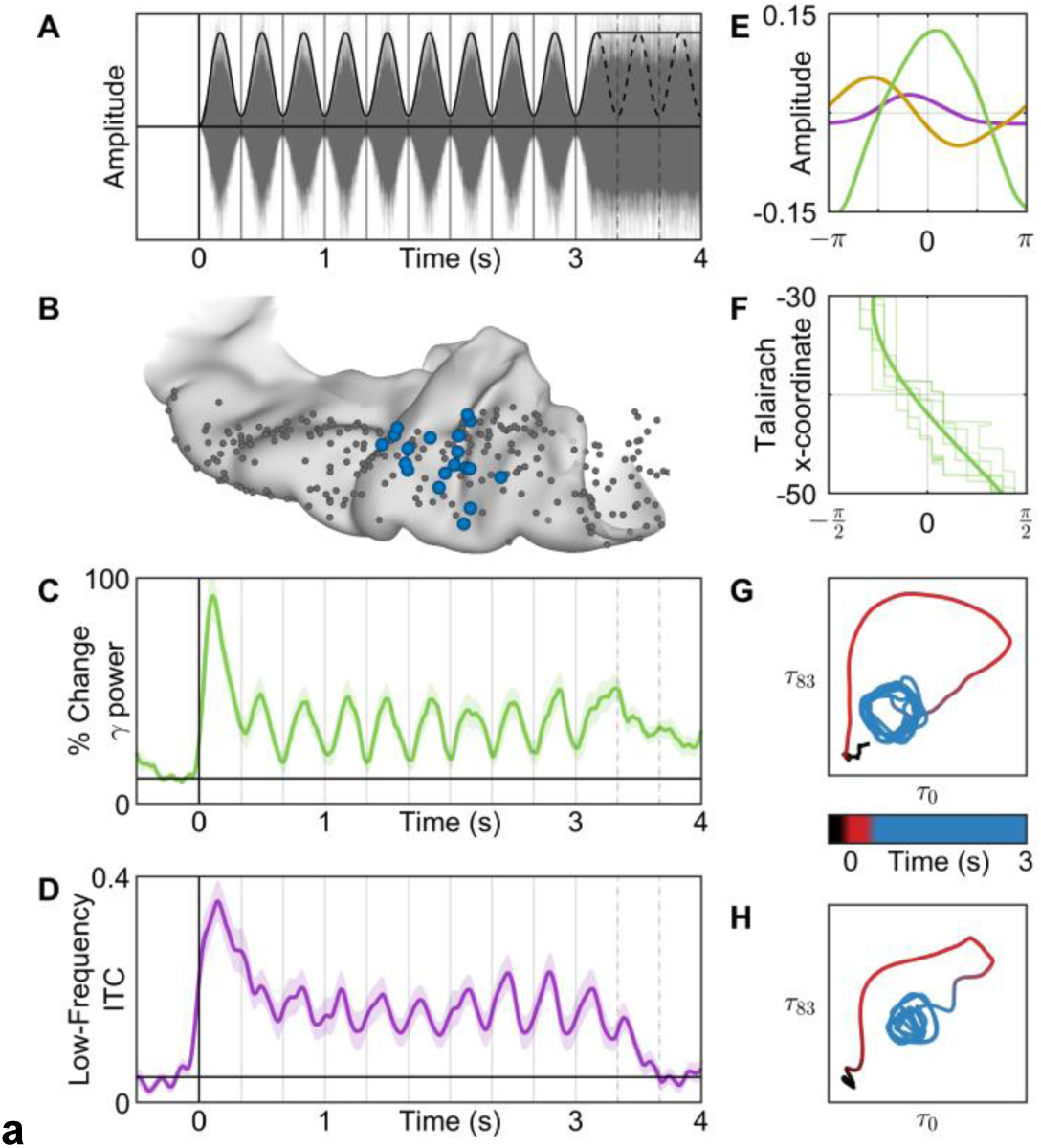
Cortical Entrainment to Rhythmic White Noise. Cortical response to rhythmic white noise. (**A**) 3 Hz amplitude-modulated white noise stimulus. (**B**) The most entrained electrode (blue) was selected from all electrodes (grey) in each patient with a supratemporal depth probe (n = 21). These were used for the following analyses. (**C**) Average percent change in gamma (65-115 Hz) power relative to pre-stimulus baseline. (**D**) Average absolute change in low-frequency (2-15 Hz) inter-trial coherence (ITC) from a pre-stimulus baseline. (**E**) Average amplitude of low (purple), beta (yellow), and gamma (green) frequencies relative to stimulus phase during entrainment (pulses 2-9) demonstrating frequency-multiplexed encoding of acoustic envelope. (**F**) Spatial distribution of peak gamma power timing relative to stimulus phase demonstrates a traveling wave (velocity 0.1 m/sec) that begins medially at the insular boundary (top) and progresses to the lateral edge (bottom). The mean wave (dark lines) is superimposed over the wave at each pulse (light lines). (**G, H**) Phase space trajectory at a quarter period delay (83 ms) in gamma power (top) and low-frequency ITC (bottom). Time indicated by color: pre-stimulus baseline (black), red (first acoustic pulse), blue (pulses 2-9). Online video (**I**): Surface-based mixed-effects multilevel analysis of gamma power at all electrodes in superior temporal gyrus during listening to this 3 Hz amplitude-modulated white noise stimulus.

Gamma, beta, and low-frequency power together yielded a frequency-multiplexed encoding of acoustic envelope (Figure 1E). Gamma power was in-phase with the stimulus, beta power was resynchronized at the trough of the stimulus, and low-frequency power was modulated by the rising edge of each pulse. The unique encoding at each frequency band suggests distinct functional channels for acoustic processing. In contrast, only low-frequency phase was reset at the rising slope of each pulse – the acoustic edge. Phase reset was not observed in beta or gamma. These encodings in power and phase were found to generalize for faster modulations of the temporal envelope (5 Hz and 7 Hz, Extended Data 3).

During entrainment, we also resolved the spatiotemporal topography of gamma power along the mediolateral extent of HG/TTS (Figure 1F). A traveling wave of cortical activity coincided with each acoustic pulse, beginning at medial HG/TTS adjacent to the inferior circular sulcus of the insula and propagating laterally across the supratemporal plane to the lip of the lateral fissure (see video: Figure 1I). Each wave began approximately 80 ms before the acoustic pulse maximum and ended approximately 80 ms afterwards, traversing HG/TTS at a speed of 0.1 m/s.

### Distinct Substrates Encode Acoustic Onset and Entrainment

Immediately posterior to HG/TTS in the planum temporale (PT), a distinct functional region generated a transient response to white noise. This region featured a high-magnitude increase in gamma power accompanied by broadband low-frequency phase reset that returned to pre-stimulus baseline activity after a single acoustic pulse. We separated this transient response from entrainment using non-negative matrix factorization – an unsupervised clustering algorithm – across all supratemporal electrodes (n = 289). This analysis revealed a distinct anteroposterior response gradient from sustained entrainment in HG/TTS to transient activity in PT (Figure 2B,D,G). This spatial distribution was significant for both gamma power (Figure 2A; r_s_ = 0.4392, p < 10^−5^) and low-frequency phase (Figure 2C; r_s_ = 0.7508, p < 10^−16^). Classification by both measures were strongly correlated (Figure 2E; r_s_ = 0.4768, p < 10^−8^); only 4 of 289 electrodes showed a mixed classification (i.e. entrainment bias in gamma power with transient bias in low-frequency phase, or the reverse; Figure 2F). The entrainment response was noted in both language dominant and non-dominant cortex, but the transient response was limited to language dominant cortex.

**Figure 2:**
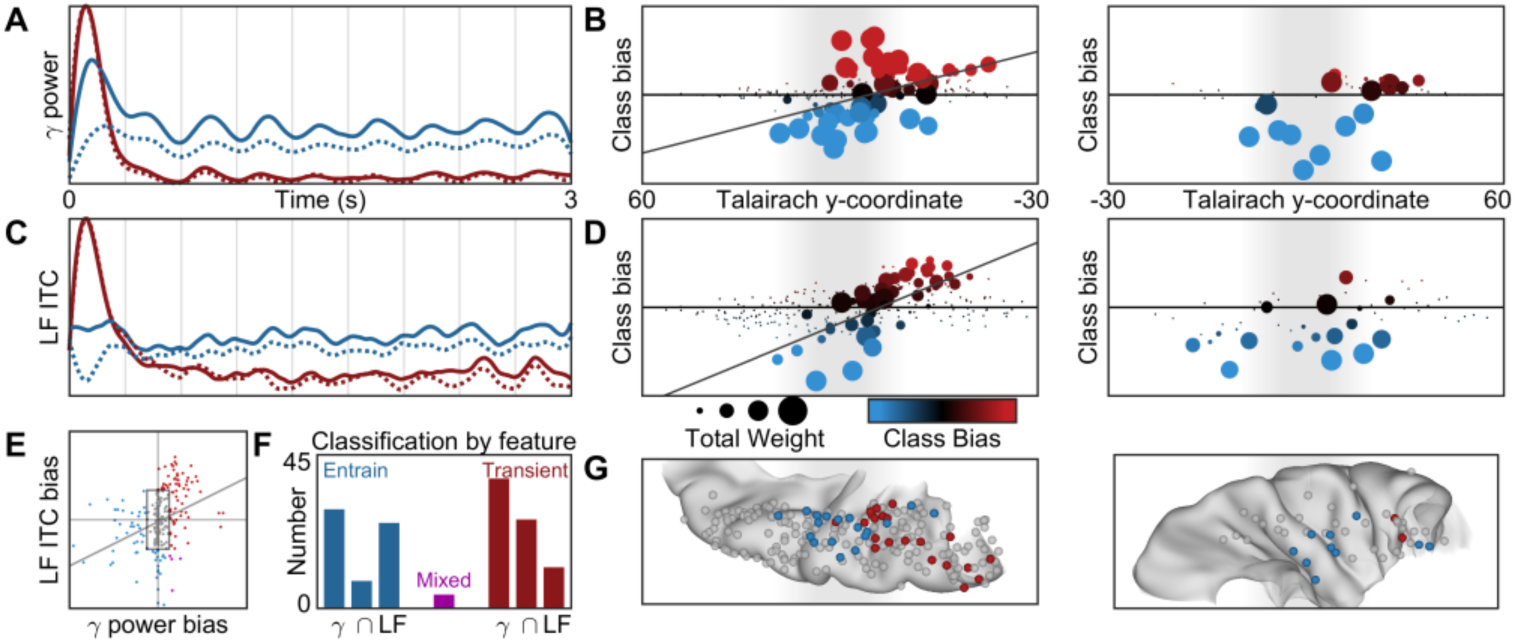
Supratemporal Distribution of Entrainment and Transient Responses. Supratemporal responses (n = 289 electrodes) classified with 2-basis non-negative matrix factorization. (**A**) Gamma power identifies an entrainment (blue) and transient (red) response: normalized basis functions (dotted line) and the normalized group-average response for the top 10% of electrodes in each class (solid line). (**B**) Spatial distribution of activation (sum of class weights; point size) and bias (difference in class weights; point color) reveals anteroposterior gradient of functional response. The left panel shows electrodes in language dominant cortex; the right, in language nondominant cortex. (**C, D**) Separately, low-frequency ITC also revealed sustained and transient responses with the same spatial distribution. (**E**) The class bias determined by gamma power and low-frequency ITC analyses were significantly correlated. (**F**) Class biases greater than a value of 5 generated discrete classifications: entrainment (n = 62), transient (n =76), or mixed (n = 4). (**G**) Electrode classifications are shown on a standard supratemporal atlas, demonstrating a clear functional split between Heschl’s gyrus and planum temporale in language dominant cortex.

The spatial topology of early auditory cortical responses was further elucidated within a single patient who underwent two separate implants (Extended Data 1): one with surface grid and another with depth electrodes. Strong entrainment encoding in HG/TTS and a robust transient response in PT were observed at electrodes along the supratemporal depth probe, but not at *any* subdural electrodes directly overlying superior temporal gyrus. This unique case indicates that the entrainment and transient responses are selectively encoded by early auditory cortex. Crucially, lateral superior temporal gyrus does not appear to be engaged in acoustic entrainment driven by sublexical features^21^.

### Prediction of Low-Level Acoustic Stimuli

To isolate neural mechanisms supporting prediction in HG/TTS, we quantified the persistence of entrainment to a 3 Hz acoustic envelope after the stimulus rhythm ceased (Figure 3A). Low-frequency phase maintained an entrained state for one cycle after the last acoustic pulse (Figure 3B); by the second cycle, this temporal organization of cortical phase was not significantly distinct from pre-stimulus baseline. In contrast, the entrained relationship between gamma power and the acoustic envelope did not carry predictive information in either cycle after the last acoustic pulse (Figure 3C). Thus, prediction in early auditory cortex is best modeled by low-frequency phase reset at acoustic edges. This neural mechanism is engaged within a single cycle of a rhythmic acoustic stimulus and remains active for at least one cycle afterwards. Such a neuro-computational solution for entrainment and prediction provides a neurobiological basis for cognitive models of speech perception^4,8,9,22^.

**Figure 3:**
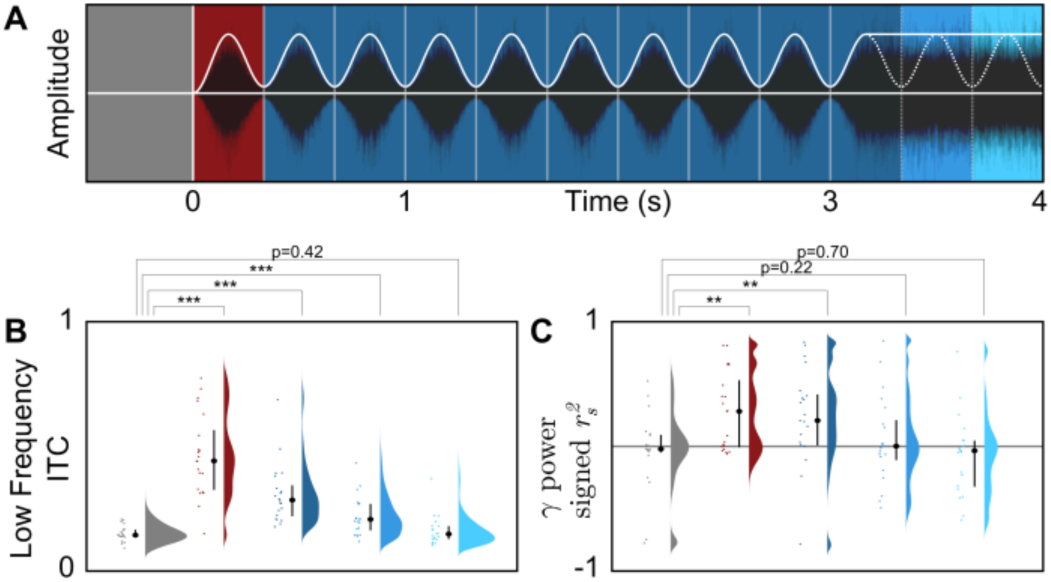
Prediction in Entrained Auditory Cortex. Low-frequency phase in early auditory cortex shows evidence of predictive encoding. (**A**) The stimulus was divided into 5 intervals: baseline (grey), onset (red), entrainment (dark blue), early prediction (medium blue), and late prediction (light blue). Crucially, there is no modulation of white noise amplitude in either of the prediction intervals. The same electrode group shown in Figure 1B was used for the following analyses. (**B, C**) The degree of entrainment in low-frequency phase and gamma power during each interval (* p < 0.05, ** p < 0.01, *** p < 0.001). Entrainment in low-frequency phase was measured as average ITC; entrainment in gamma power was measured as a signed *r*^2^ from the Spearman’s correlation with a 3 Hz sine wave. Both low-frequency phase and gamma power were significantly entrained during the onset and entrainment intervals, but only low-frequency phase remained significantly entrained during the first prediction interval.

### Entrainment to Natural Language Speech

In a second experiment, patients (n = 20) named common objects cued by short spoken descriptions (e.g. they heard “a place with sand along a shore” and articulated “beach”). For each sentence (Figure 4A), we extracted a pair of key features suggested by our analysis of rhythmic white noise: acoustic envelope and edges. The former describes the instantaneous amplitude of speech, while the latter demarcates moments of rapid amplitude gain. We evaluated the engagement of neural substrates that entrained to the white noise stimulus during natural language speech. Power in HG/TTS was significantly correlated with the acoustic envelope of speech (low-frequency, r_s_ = −0.0659, p < 10^−3^; beta, r_s_ = −0.0532, p < 10^−3^; gamma, rs = 0.0736, p < 10^−3^; Figure 4B) at a frequency-specific delay (low-frequency, 135 ms; beta, 100 ms; gamma, 60 ms; Figure 4C). Low-frequency phase organization in HG/TTS was significantly increased during the 125 ms following acoustic edges in speech (p < 10^−3^; Figure 4D). Furthermore, it was significantly greater following acoustic edges than following syllabic onsets (p = 0.0061; Figure 4D) – a similar characteristic, but derived from and specific to speech. These findings are concordant with the frequency-multiplexed encoding of acoustic envelope and the low-frequency phase reset at acoustic edges observed during entrainment to the white noise stimulus. The neural response was preserved during reversed speech, emphasizing the sublexical nature of this process. The cortical encoding of the speech envelope (Figure 4C) and of edges (Figure 4E) was localized to HG/TTS – the same supratemporal region that demonstrated entrainment and prediction for the white noise stimulus.

**Figure 4:**
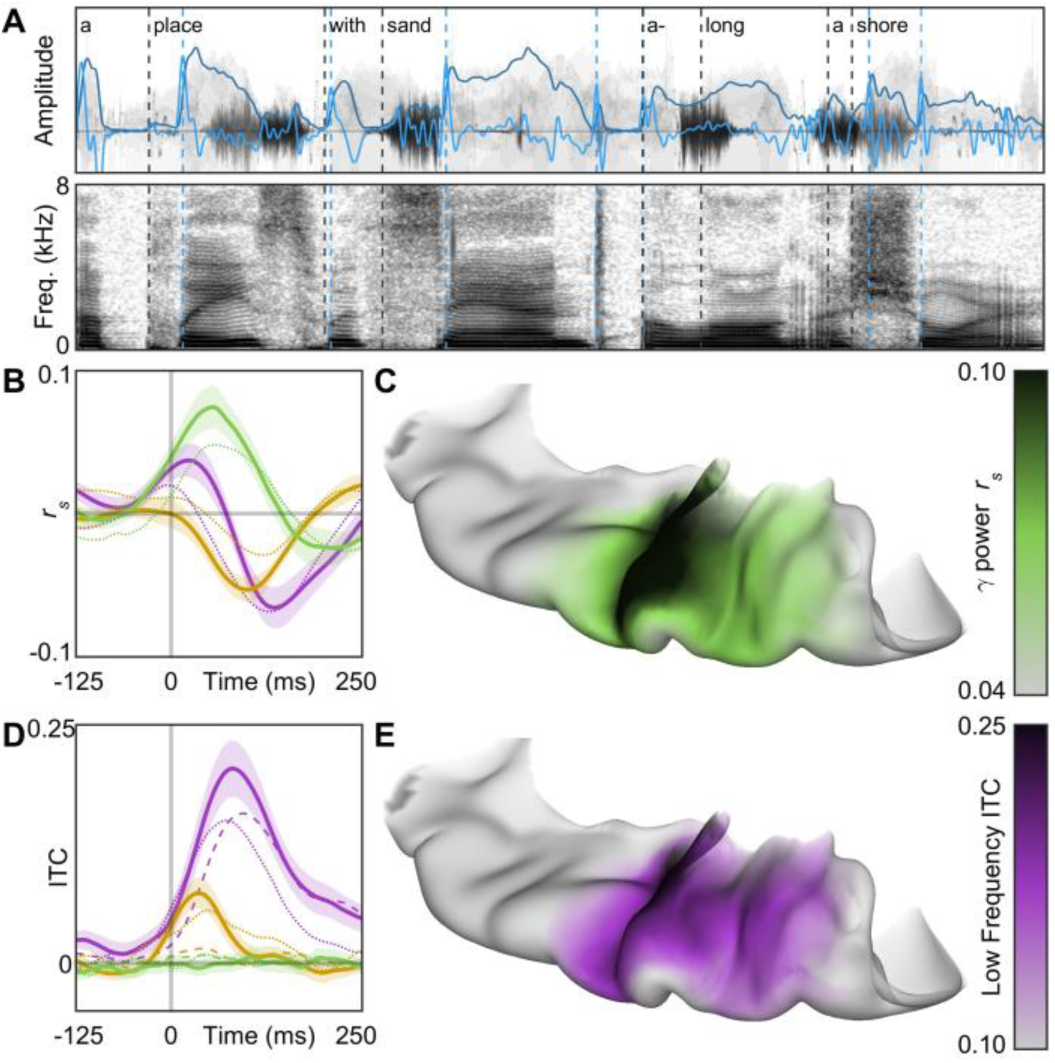
Cortical Entrainment to Speech. Entrainment to natural language speech occurs focally in early auditory cortex. (**A**) Patients listened to short sentences describing common objects. Two features were extracted from each stimulus: acoustic envelope (light blue) and acoustic edges (dark blue). For comparison, syllabic onsets (black) were also demarcated. (**B**) The peak lagged Spearman’s correlation between acoustic and gamma envelopes and (**D**) the average low-frequency ITC following an acoustic edge were computed for all electrodes in superior temporal gyrus. These measures were mapped onto a standard MNI atlas, revealing acoustic encodings limited to early auditory cortex. (**C**) The encoding of acoustic envelope for speech perception was further explored by cross correlation with low-frequency (purple), beta (yellow), and gamma (green) amplitudes. The effect size was slightly reduced for reversed speech (dotted lines). (**E**) Acoustic edges were better encoded by low-frequency phase than syllabic onsets (dashed lines). Online video (**F**): Surface-based mixed-effects multilevel analysis of gamma power at all electrodes in superior temporal gyrus during listening to natural language speech.

Natural language speech recruited a much broader set of neuroanatomic substrates than white noise, including planum polare, lateral superior temporal gyrus, and superior temporal sulcus (Figure 4F). In the patient with both surface grid and depth electrodes, only speech induced significant activity in the lateral temporal grid electrodes (Extended Data 1). This supplementary speech-specific cortex is presumably engaged for the downstream processing of higher-order language features (e.g. phonemes^23^).

### Supratemporal Dissociations in Speech Perception and Production

We compared neural activity in both HG/TTS and PT during listening and speaking – externally and internally generated speech. In each patient with a supratemporal depth probe, the pair of electrodes with the strongest entrainment and transient responses were identified during the rhythmic white noise condition. These criteria selected electrodes in HG/TTS and PT, respectively (Figure 5A). Gamma power in these regions was analyzed relative to sentence and articulation onset for a representative individual (Figure 5B) and across the group (Figure 5C). HG/TTS responded strongly during both listening and speaking, remaining active for the duration of each sentence and throughout articulation. In contrast, PT also responded strongly following sentence onset with peak activity at 100 ms; however, this region was quiescent during articulation (Figure 5D).

**Figure 5:**
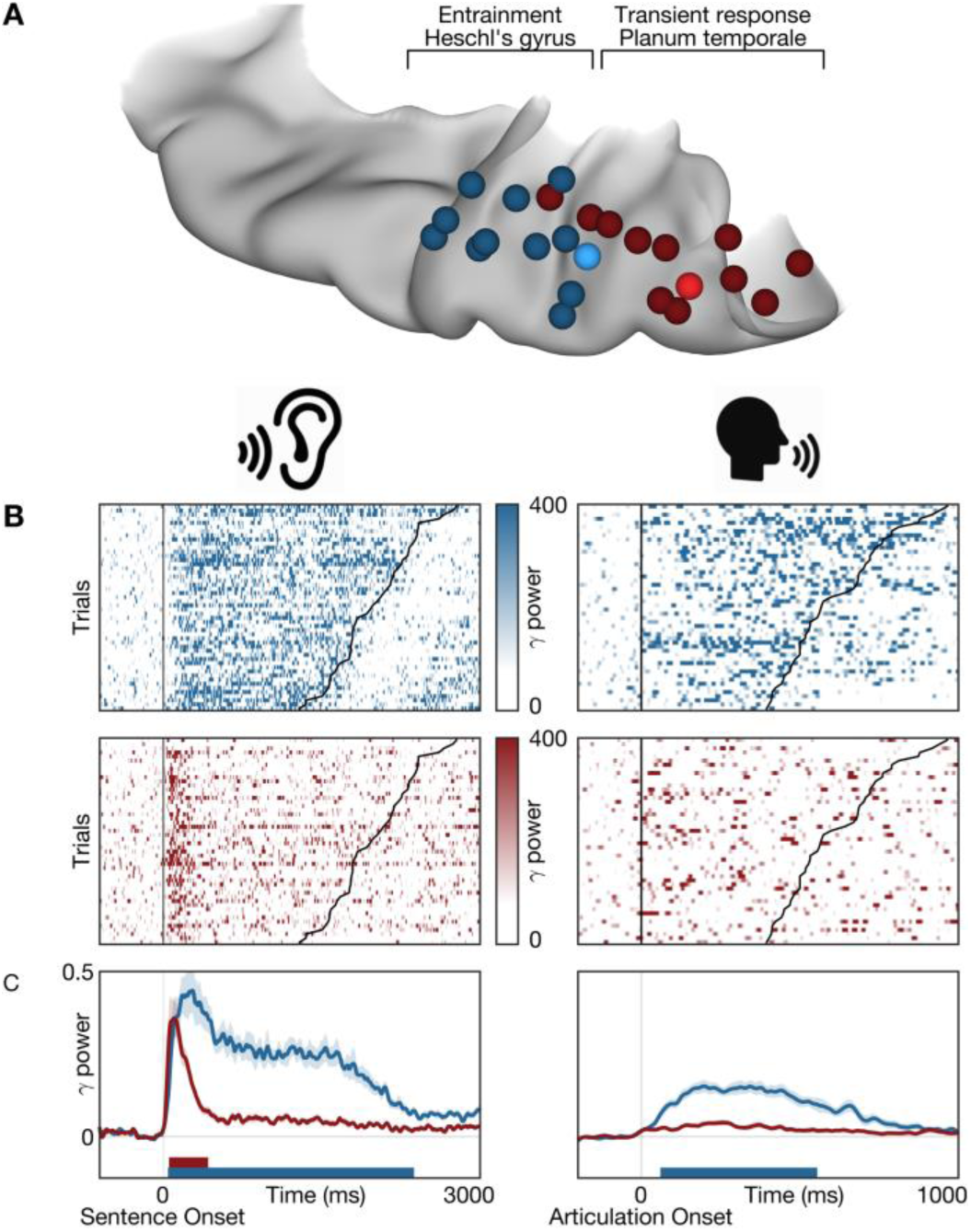
Distinct Supratemporal Responses During Listening and Speaking. Functional dissociation in HG/TTS and PT during listening and speaking. (**A**) A pair of electrodes were selected in each patient for an entrainment (blue) and transient (red) response to the 3 Hz amplitude-modulated white noise stimulus. One representative patient was highlighted (bright electrode pair) for single-trial analysis. (**B**) Single-trial raster plots of the percent change in gamma power during speech listening and production. (**C**) Gamma power averaged across trials and then across patients. Significance bars were determined at an alpha level of p < 0.001 with familywise error correction. While the gamma response in HG/TTS is reduced during articulation, the response in PT is eliminated. Online video (**D**): Surface-based mixed-effects multilevel analysis of gamma power at all electrodes in superior temporal gyrus during single-word articulations.

We further characterized the spatial distribution of the transient response during speech listening and its suppression during speech production using non-negative matrix factorization. As for the white noise stimulus, gamma power yielded sustained entrainment and transient response types (Figure 6A) along a robust anteroposterior distribution (Figure 6E,G; r_s_ = 0.4869, p < 10^−10^). These were strongly correlated with the class biases for white noise listening (Figure 6B; r_s_ = 0.6029, p < 10^−20^). When this factorization was applied to gamma power during articulation (Figure 6C,F,H), the sustained entrainment response was preserved (r_s_ = 0.6764, p < 10^−16^) while the transient response type was suppressed (r_s_ = 0.0935, p = 0.4726). Of the 30 electrodes demonstrating a transient response during speech listening, only 1 retained this classification during articulation (Figure 6D). The functional dissociation at PT between externally and internally generated speech provides the first direct evidence for the theory of predictive coding during speech production^22,24^ via motor-to-sensory feedback^25–27^.

**Figure 6:**
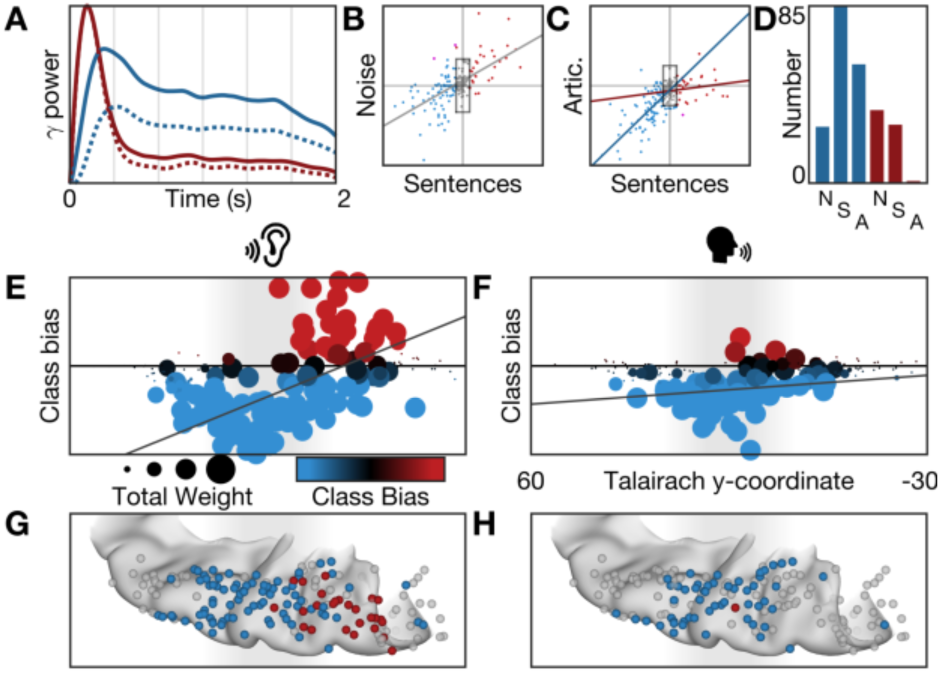
Transient Response is Uniquely Suppressed for Self-Generated Speech. Supratemporal responses (n = 188 electrodes) classified with 2-basis non-negative matrix factorization. (**A**) Gamma power identifies an entrainment (blue) and transient (red) response: normalized basis functions (dotted line) and the normalized group-average response for the top 10% of electrodes in each class (solid line). (**B**) The class bias determined by factorizations of electrode responses to noise and sentence listening – homogenous and structure acoustic inputs – were significantly correlated. (**C**) The factorization from sentence listening was applied to the electrode responses at articulation. Entrainment class bias was significantly correlated for listening and speaking, but the transient class bias was uncorrelated. (**D**) Class biases greater than a value of 10 generated discrete classifications for speech listening (entrainment, n = 85; transient, n = 29) and articulation (entrainment, n = 57; transient, n = 1). (**E**, **F**) Spatial distribution of activation (sum of class weights; point size) and bias (difference in class weights; point color) reveals anteroposterior gradient of functional response during speech listening (left) but not articulation (right). (**G**, **H**) Electrodes shown on a standard supratemporal atlas reveal that the transient response is localized to PT and is uniquely suppressed during articulation.

## Discussion

Direct intracranial recordings of the supratemporal plane resolved the functional architecture of entrainment and prediction in human early auditory cortex at an unprecedented resolution and scale. Entrainment to speech engages a frequency-multiplexed encoding of two sublexical acoustic features: envelope and edge. A pair of distinct neuroanatomic substrates perform predictive encoding: “when” by HG/TTS in low-frequency phase and “what” by PT in gamma power. The identification and characterization of these mechanisms advances the understanding of how human cerebral cortex parses continuous acoustic input during both speech perception and production.

### Entrainment and Prediction of “When” in Heschl’s Gyrus

Entrainment is the synchronization of two quasi-periodic systems – intrinsic neural oscillations with extrinsic rhythmic signals. It is thought to play an important role in a variety of cognitive processes including attentional selection^7,28,29^ and internal timekeeping^30–33^. Furthermore, entrainment is axiomatic to leading models of speech comprehension^4,8,9,22^. These theories are supported by evidence that speech envelope distortions impair comprehension^34–36^ independent of spectral content^37–39^ and that the degree of neuro-acoustic entrainment modulates intelligibility^40–43^.

Entrainment has been variably characterized as either the encoding of envelope amplitude in bandlimited cortical power^44–47^ or of discrete segmental events in evoked response potentials^7,10,41,48,49^. Using electrodes positioned along the anteroposterior extent of the supratemporal plane, we localized the cortical signature of entrainment to strictly early auditory cortex: Heschl’s gyrus and the transverse temporal sulcus^50–52^. This signature was considerably more complex than that suggested by prior studies, comprising a frequency-multiplexed encoding of envelope phase – distinct for rising and falling amplitudes of the same magnitude – in low-frequency, beta, and gamma power. We also identified a separate, concurrent encoding of acoustic edges in low frequency phase reset. This encoding uniquely persisted after the entraining stimulus ended, consistent with the behavior of a predictive neural mechanism. Importantly, identical cortical substrates of entrainment were engaged during natural sentence listening.

Our findings generate insights to the nature of entrainment and its support of speech perception. First, acoustic entrainment has been described by others as either a “continuous mode” for acoustic processing^29,53–57^ or simply a recurring series of transient evoked responses^10,58,59^. The former interpretation is most consistent with our observations of a non-adapting entrained state that is distinct from the evoked response at stimulus onset and that endures after the entraining stimulus ends. Second, in contrast to prior studies^42,45^, we found that reversed speech drove an equivalent degree of entrainment in early auditory cortex; furthermore, acoustic edges were more strongly encoded in cortical phase than syllabic onsets – a linguistic feature with similar frequency and periodicity. This supports the assertion^60^ that entrainment is driven by sublexical acoustic processing, perhaps even inherited from subcortical regions (e.g. medial geniculate nucleus). Third, the multiband encoding of acoustic envelope in cortical power is richer than has been previously suggested^4,8,9^. While gamma power does track the instantaneous acoustic envelope, both beta and low frequency power contribute unique information. This supports frequency-multiplexed acoustic processing^22,61–63^ with each band representing distinct channels of information exchange^64–67^.

The utility of entrainment is thought to be the organization of transient excitability states within neuronal populations^68–72^. Discrete high excitability periods constitute “windows of opportunity” for input into sensory cortex, as evidenced by peri-threshold detection studies in somatosensory^73^, visual^33,74,75^, and auditory^46,60,76^ regions. During listening, such windows might serve to segment speech to facilitate comprehension^77^. More generally, the temporal organization of high excitability periods could serve to minimize temporal uncertainty in stimulus processing and detection^7,78–80^. This view was corroborated by a behavioral study of responses to the same white noise stimulus used in these experiments that revealed a striking relationship between detection accuracy and the preceding rhythmic stimulus^81^. With direct intracranial recordings, we found that low-frequency phase reset anticipates the first “missing” acoustic edge.

These results constitute strong evidence for neural mechanisms in early auditory cortex supporting entrainment and prediction, both fundamental computational elements in models of speech perception^4,8,9,22^. The characteristics of these mechanisms contrast with the presumption of “a principled relation between the time scales present in speech and the time constants underlying neuronal cortical oscillations that is both a reflection of and the means by which the brain converts speech rhythms into linguistic segments”^4^ supported by “cascaded cortical oscillations”^8^ or a “hierarchy of nested oscillations”^9^. Our results are instead consistent with the predictive encoding of “when” by a bandlimited complex of discrete computational channels, each arising from distinct patterns of hierarchical cortical connectivity^22^.

### Transient Response and Prediction of “What” in Planum Temporale

While entrainment was constrained to Heschl’s gyrus and the transverse temporal sulcus, we observed a distinct transient response in planum temporale. The transient response was characterized by a brief spike in gamma power and rapid reset of low-frequency phase immediately following acoustic onset. Interestingly, this response was not engaged during self-generated speech. Such preferential engagement for unexpected sound is consistent with predictive encoding during speech production^26,27^. Upon execution of a speech motor plan, a learned internal model generates an efference copy^82–84^ – an expected sensory result. When the acoustic input matches this efference copy, no cortical signal is generated; however, when a mismatch occurs (e.g. externally-generated sound or speech), an error signal results^22^. This is precisely what we observed in the planum temporale, distinct from entrainment in Heschl’s gyrus.

Our results advance understanding of the neurobiology of predictive speech coding in two respects. First, functional studies have revealed single-unit preference in primary auditory cortex for listening or speaking in both non-humans^85^ and humans^82^. It has recently been asserted that these response tunings overlap – an “intertwined mosaic of neuronal populations”^86^ in auditory cortex. Instead, the complete anteroposterior mapping of the supratemporal plane in 31 patients enabled us to identify a distinct neuroanatomical organization in planum temporale. Second, several groups report cortical response suppression specific for self-generated speech^19,86–89^. We reveal two distinct modes that enable this suppression: a partial reduction of activity in Heschl’s gyrus and a complete absence of the transient response in planum temporale. The stapedius reflex^85,90^ does not explain the latter mode, suggesting a neural mechanism of suppression. All together, we provide compelling evidence for efference copies – predictive encoding of “what”^22^ – and their essential role in speech production^26,27^.

## Acknowledgements

We thank all the patients who participated in this study; laboratory members at the Tandon lab (Matthew Rollo and Jessica Johnson); neurologists at the Texas Comprehensive Epilepsy Program (Jeremy Slater, Giridhar Kalamangalam, Omotola Hope, Melissa Thomas) who participated in the care of these patients; and all the nurses and technicians in the Epilepsy Monitoring Unit at Memorial Hermann Hospital who helped make this research possible.

This work was supported by the National Institute on Deafness and Other Communication Disorders 5R01DC014589-04, National Institute of Neurological Disorders and Stroke 5U01NS098981-03, and National Institute on Deafness and Other Communication Disorders 1F30DC017083-01.

## Author Contributions

Conceptualization: GH, NT; Methodology: KJF, GH, NT; Software: KJF, PSR; Formal Analysis: KJF; Investigation: KJF; Data Curation: KJF, PSR, NT; Writing – Original Draft: KJF; Writing – Review & Editing: KJF, GH, NT; Visualization: KJF; Supervision: NT; Project Administration: NT; Funding Acquisition: NT.

## Competing Interests

The authors declare no competing interests.

## Materials & Correspondence

Please direct all correspondence and material requests to Nitin Tandon.

## Methods

### Population

31 patients (18 male, 13 female; mean age 31 ± 8; mean IQ 96 ± 15) undergoing evaluation of intractable epilepsy with intracranial electrodes were enrolled in the study after obtaining informed consent. Study design was approved by the committee for the protection of human subjects at the University of Texas Health Science Center. A total of 6580 electrodes (5742 depths, 838 grids) were implanted in this cohort. Only the 4003 electrodes (3494 depths, 509 grids) unaffected by epileptic activity, artifacts, or electrical noise were used in subsequent analyses.

Hemispheric language dominance was evaluated in all patients with intra-carotid sodium amytal injection^91^ (n = 5), fMRI laterality index^92,93^ (n = 7), cortical stimulation mapping^94^ (n = 8), or the Edinburgh Handedness Inventory^95^ (n = 11). 29 patients were confirmed to be left-hemisphere language-dominant. 1 patient was found to be left-handed by EHI and did not undergo alternative evaluation; they are assumed to be left-hemisphere dominant, but were excluded from laterality analysis. Three patients were found to be right-hemisphere language-dominant; 2 by intra-carotid sodium amytal injection and 1 by fMRI laterality index.

### Paradigms

Two distinct paradigms were used. The first experiment featured amplitude-modulated white noise, while the latter experiment contained natural speech. All were designed to evaluate the response of early auditory cortex to external acoustic stimuli. Stimuli were played to patients using stereo speakers (44.1 kHz, 15” MacBook Pro 2013) driven by either MATLAB (first experiment) or Python (second experiment) presentation software.

The first experiment presented patients with a single-interval two-alternative forced-choice perceptual discrimination task^81^. The stimulus comprised two periods. In the first, wideband Gaussian noise was modulated (3 Hz, 80% depth) for 3 seconds. In the second, the modulation waveform ended on the cosine phase of the next cycle to yield 833 ms of constant-amplitude noise. Furthermore, 50% of trials featured a peri-threshold tone (1 kHz, 50 ms duration, 5ms rise-decay time) that was presented at one of 6 temporal positions and at an amplitude level from 1 of 3 values. The temporal positions were separated by a quarter-cycle of the modulation frequency beginning with the constant-amplitude noise. The amplitude levels covered a range of 12 dB. On each trial, the patient was required to indicate via a key press whether a tonal signal was present during the unmodulated segment of the masking noise. All patients each completed 100 trials.

In the second experiment, patients engaged in an auditory-cued naming task: naming to definition. The stimuli were single sentence descriptions (average duration of 1.97 ± 0.36 seconds) recorded by both male and female speakers. These were designed such that the last word always contained crucial semantic information without which a specific response could not be generated (e.g., “A round red *fruit*.”)^96^. Patients were instructed to articulate aloud the object described by the stimulus. In addition, temporally-reversed speech was used as a control condition. These stimuli preserved the spectral content of natural speech, but communicated no meaningful linguistic content. For each stimulus, patients were instructed to articulate aloud the gender of the speaker. 20 patients each completed 180 trials.

### MR acquisition

Pre-operative anatomical MRI scans were obtained using a 3T whole-body MR scanner (Philips Medical Systems) fitted with a 16-channel SENSE head coil. Images were collected using a magnetization-prepared 180° radiofrequency pulse and rapid gradient-echo sequence with 1 mm sagittal slices and an in-plane resolution of 0.938 × 0.938 mm^97^. Pial surface reconstructions were computed with FreeSurfer (v5.1)^98^ and imported to AFNI^99^. Post-operative CT scans were registered to the pre-operative MRI scans to localize electrodes relative to cortical landmarks. Grid electrode locations were determined by a recursive grid partitioning technique and then optimized using intra-operative photographs^100^. Depth electrode locations were informed by implantation trajectories from the ROSA surgical system.

### ECoG acquisition

Stereo-electroencephalographic depth probes with platinum-iridium electrode contacts (PMT Corporation; 0.8 mm diameter, 2.0 mm length cylinders; adjacent contacts separated by 1.5-2.43 mm) were implanted using the Robotic Surgical Assistant (ROSA; Zimmer-Biomet, Warsaw, IN) registered to the patient using both a computed tomographic angiogram and an anatomical MRI^101,102^. Each depth probe had 8-16 contacts and each patient had multiple (12-16) such probes implanted. Surface grids – subdural platinum-iridium electrodes embedded in a silastic sheet (PMT Corporation, Chanhassen, MN; top-hat design; 3 mm diameter cortical contact) – were surgically implanted via a craniotomy^93,100,103^. ECoG recordings were performed at least two days after the craniotomy to allow for recovery from the anesthesia and narcotic medications. 29 patients were implanted with depth probe electrodes; 4 patients were implanted with surface grid electrodes. Notably, a pair of patients had 2 separate implants: first with depth probe electrodes and subsequently with surface grid electrodes.

Data were collected at a 2000 Hz sampling rate and 0.1-700 Hz bandwidth using NeuroPort NSP (Blackrock Microsystems, Salt Lake City, UT). Stimulus presentation software triggered a digital pulse at trial onset that was registered to ECoG via digital-to-analog conversion (MATLAB: USB-1208FS, Measurement Computing, Norton, MA; Python: U3-LV, LabJack, Lakewood, CO). Continuous audio registered to ECoG was recorded with an omnidirectional microphone (30-20,000 Hz response, 73 dB SNR, Audio Technica U841A) placed adjacent to the presentation laptop. For the naming to definition and reversed speech experiments, articulation onset and offset were determined by offline analysis of the amplitude increase and spectrographic signature associated with each verbal response.

Cortical areas with potentially abnormal physiology were excluded by removing channels that demonstrated inter-ictal activity or that recorded in proximity to the localized seizure onset sites. Additional channels contaminated by >10 dB of line noise or regular saturation were also excluded from further analysis. The remaining channels were referenced to a common average comprised of all electrodes surviving these criteria. Any trials manifesting epileptiform activity were removed. Furthermore, trials for the naming to definition and reversed speech experiments in which the patient answered incorrectly or after more than 2 seconds were eliminated.

### Digital signal processing

Line noise was removed with zero-phase 2^nd^ order Butterworth bandstop filters at 60 Hz and its first 2 harmonics. The analytic signal was generated with frequency domain bandpass Hilbert filters featuring paired sigmoid flanks (half-width 1 Hz)^104–107^. For spectral decompositions, this was generalized to a filter bank with logarithmically spaced center frequency (2 to 16 Hz, 50 steps) and passband widths (1 to 4 Hz, 50 steps). Instantaneous amplitude and phase were subsequently extracted from the analytic signal. In this fashion, we used both narrowband and wideband analyses to precisely quantify the frequency driving a local cortical response and its timing, respectively.

### Statistical analysis

Analyses were performed with trials time-locked to stimulus onset (all experiments) and to articulation onset (only naming to definition and reversed speech experiments). The baseline period for all experiments was defined as −300 to −50 ms relative to stimulus onset.

Instantaneous amplitude was squared and then normalized to the baseline period, yielding percent change in power from baseline. Statistical significance was set at the 0.01 level, evaluated with the Wilcoxon signed-rank test, and subjected to familywise error correction. The alignment of instantaneous phase at each trial *θ*_*n*_was quantified with inter-trial coherence (ITC), defined as follows where *N* is the number of trials:

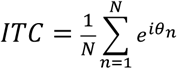

Statistical significance was set at the 0.01 level, evaluated with the Rayleigh *z* test, and subjected to FDR control. All time traces were smoothed after statistical analysis with a Savitsky-Golay polynomial filter (3^rd^ order, 83ms frame length) for visual presentation.

Non-negative matrix factorization (NNMF) is an unsupervised clustering algorithm^108^. This method expresses non-negative matrix **A** ∈ *R*^mxn^ as the product of “class weight” matrix **W** ∈ *R*^mxk^ and “class archetype” matrix **H** ∈ *R*^kxn^, minimizing:

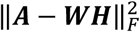

The factorization rank *k* = 2 was chosen for all analyses in this work. Repeat analyses with higher ranks did not identify additional response types. We optimized the matrix factorization with 1000 replicates of a multiplicative update algorithm (MATLAB R2018b Statistics and Machine Learning Toolbox). Two types of inputs were separately factorized: mean gamma power and low-frequency phase ITC. Gamma power values less than 20% above baseline and low-frequency phase ITC less than baseline were rectified. These were calculated for the *m* electrodes in the supratemporal plane at *n* time points. Factorization thus generated a pair of class weights for each electrode and a pair of class archetypes – the basis function for each class. Class bias was defined as the difference between the class weights at each electrode. Response magnitude was defined as the sum of class weight magnitudes at each electrode. Separate factorizations were estimated for white noise listening and for natural speech listening. The latter was then applied to self-generated speech by conserving the class archetypes and recalculating the class weights:

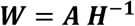

Response classifications were established by applying a binary threshold to the class biases. Spearman correlations were calculated using only electrodes with a large response magnitude (>20).

Surface-based mixed-effects multilevel analysis (SB-MEMA) was used to provide statistically robust^109–111^ and topologically precise^107,112,113^ effect estimates of band-limited power change from the baseline period. This method, developed and described previously by our group^105,114^, accounts for sparse sampling, outlier inferences, as well as intra- and inter-subject variability to produce population maps of cortical activity. SB-MEMA was run on short, overlapping time windows (150 ms width, 10 ms spacing) to generate the frames of a movie portraying cortical activity. All maps were smoothed with a geodesic Gaussian smoothing filter (3 mm full-width at half-maximum) for visual presentation.

## Data Availability

The datasets collected and analyzed during the current study are not publicly available as patients did not consent to such distribution but grouped data representations are available from the corresponding author on reasonable request.

## References

1. Heald, S., Klos, S. & Nusbaum, H. in Neurobiology of Language (2016). doi:10.1016/B978-0-12-407794-2.00017-1

2. Lotto, A. J. & Holt, L. L. Speech perception: The view from the auditory system. Neurobiology of Language (2016). doi:10.1016/B978-0-12-407794-2.00016-X

3. Rosen, S. Temporal information in speech: acoustic, auditory and linguistic aspects. Philosophical transactions of the Royal Society of London. Series B, Biological sciences 336, 367–373 (1992).

4. Giraud, A.-L. L. & Poeppel, D. Cortical oscillations and speech processing: Emerging computational principles and operations. Nat. Neurosci. 15, 511–7 (2012).

5. Obleser, J., Henry, M. J. & Lakatos, P. What do we talk about when we talk about rhythm? PLoS Biol. (2017). doi:10.1371/journal.pbio.2002794

6. Rimmele, J. M., Morillon, B., Poeppel, D. & Arnal, L. H. Proactive Sensing of Periodic and Aperiodic Auditory Patterns. Trends in Cognitive Sciences (2018). doi:10.1016/j.tics.2018.08.003

7. Schroeder, C. E. & Lakatos, P. Low-frequency neuronal oscillations as instruments of sensory selection. Trends Neurosci. 32, 9–18 (2009).

8. Ghitza, O. Linking speech perception and neurophysiology: Speech decoding guided by cascaded oscillators locked to the input rhythm. Front. Psychol. 2, (2011).

9. Peelle, J. E. & Davis, M. H. Neural oscillations carry speech rhythm through to comprehension. Frontiers in Psychology 3, (2012).

10. Doelling, K. B., Arnal, L. H., Ghitza, O. & Poeppel, D. Acoustic landmarks drive delta-theta oscillations to enable speech comprehension by facilitating perceptual parsing. Neuroimage 85, 761–768 (2014).

11. Howard, M. F. & Poeppel, D. The neuromagnetic response to spoken sentences: Co-modulation of theta band amplitude and phase. Neuroimage (2012). doi:10.1016/j.neuroimage.2012.02.028

12. Lakatos, P., Karmos, G., Mehta, A. D., Ulbert, I. & Schroeder, C. E. Entrainment of neuronal oscillations as a mechanism of attentional selection. Science (80-. ). (2008). doi:10.1126/science.1154735

13. Lakatos, P. An Oscillatory Hierarchy Controlling Neuronal Excitability and Stimulus Processing in the Auditory Cortex. J. Neurophysiol. (2005). doi:10.1152/jn.00263.2005

14. Luo, H., Liu, Z. & Poeppel, D. Auditory cortex tracks both auditory and visual stimulus dynamics using low-frequency neuronal phase modulation. PLoS Biol. (2010). doi:10.1371/journal.pbio.1000445

15. Aliu, S. O., Houde, J. F. & Nagarajan, S. S. Motor-induced suppression of the auditory cortex. J. Cogn. Neurosci. (2009). doi:10.1162/jocn.2009.21055

16. Hickok, G. Computational neuroanatomy of speech production. Nature Reviews Neuroscience (2012). doi:10.1038/nrn3158

17. Hickok, G., Houde, J. & Rong, F. Sensorimotor Integration in Speech Processing: Computational Basis and Neural Organization. Neuron (2011). doi:10.1016/j.neuron.2011.01.019

18. Houde, J. F. & Nagarajan, S. S. Speech Production as State Feedback Control. Front. Hum. Neurosci. (2011). doi:10.3389/fnhum.2011.00082

19. Houde, J. F., Nagarajan, S. S., Sekihara, K. & Merzenich, M. M. Modulation of the auditory cortex during speech: An MEG study. J. Cogn. Neurosci. (2002). doi:10.1162/089892902760807140

20. Okada, K., Matchin, W. & Hickok, G. Neural evidence for predictive coding in auditory cortex during speech production. Psychon. Bull. Rev. (2017). doi:10.3758/s13423-017-1284-x

21. Hamilton, L. S., Edwards, E. & Chang, E. F. A Spatial Map of Onset and Sustained Responses to Speech in the Human Superior Temporal Gyrus. Curr. Biol. (2018). doi:10.1016/j.cub.2018.04.033

22. Arnal, L. H. & Giraud, A. L. Cortical oscillations and sensory predictions. Trends in Cognitive Sciences 16, 390–398 (2012).

23. Hickok, G. & Poeppel, D. Dorsal and ventral streams: A framework for understanding aspects of the functional anatomy of language. Cognition 92, 67–99 (2004).

24. Shadmehr, R., Smith, M. A. & Krakauer, J. W. Error Correction, Sensory Prediction, and Adaptation in Motor Control. Annu. Rev. Neurosci. (2010). doi:10.1146/annurev-neuro-060909-153135

25. Sperry, R. W. Neural basis of the spontaneous optokinetic response produced by visual inversion. J. Comp. Physiol. Psychol. (1950). doi:10.1037/h0055479

26. Hickok, G. Computational neuroanatomy of speech production. Nat. Rev. Neurosci. (2012). doi:10.1038/nrn3158

27. Guenther, F. H. Cortical interactions underlying the production of speech sounds. J. Commun. Disord. 39, 350–365 (2006).

28. Lakatos, P., Karmos, G., Mehta, A. D., Ulbert, I. & Schroeder, C. E. Entrainment of Neuronal Attentional Selection. Science (80-. ). 320, 23–25 (2008).

29. Schroeder, C. E., Wilson, D. A., Radman, T., Scharfman, H. & Lakatos, P. Dynamics of Active Sensing and perceptual selection. Current Opinion in Neurobiology 20, 172–176 (2010).

30. Stefanics, G. et al. Phase entrainment of human delta oscillations can mediate the effects of expectation on reaction speed. J. Neurosci. 30, 13578–85 (2010).

31. Saleh, M., Reimer, J., Penn, R., Ojakangas, C. L. & Hatsopoulos, N. G. Fast and slow oscillations in human primary motor cortex predict oncoming behaviorally relevant cues. Neuron 65, 461–471 (2010).

32. Cohen, M. X. It’s about Time. Front. Hum. Neurosci. 5, (2011).

33. Cravo, A. M., Rohenkohl, G., Wyart, V. & Nobre, A. C. Endogenous modulation of low frequency oscillations by temporal expectations. J. Neurophysiol. 106, 2964–2972 (2011).

34. Ahissar, E. & Ahissar, M. in The Auditory Cortex: A Synthesis of Human and Animal Research 295–314 (2005). doi:10.4324/9781410613066

35. Ghitza, O. & Greenberg, S. On the possible role of brain rhythms in speech perception: Intelligibility of time-compressed speech with periodic and aperiodic insertions of silence. Phonetica 66, 113–126 (2009).

36. Geiser, E. & Shattuck-Hufnagel, S. Temporal regularity in speech perception: Is regularity beneficial or deleterious? Proc. Meet. Acoust. 14, 1–10 (2012).

37. Drullman, R., Festen, J. M. & Plomp, R. Effect of temporal envelope smearing on speech reception. J. Acoust. Soc. Am. 95, 1053–1064 (1994).

38. Shannon, R. V., Zeng, F. G., Kamath, V., Wygonski, J. & Ekelid, M. Speech recognition with primarily temporal cues. Science (80-. ). 270, 303–304 (1995).

39. Smith, Z. M., Delgutte, B. & Oxenham, A. J. Chimaeric sounds reveal dichotomies in auditory perception. Nature 416, 87–90 (2002).

40. Ahissar, E. et al. Speech comprehension is correlated with temporal response patterns recorded from auditory cortex. Proc. Natl. Acad. Sci. 98, 13367–13372 (2001).

41. Luo, H. & Poeppel, D. Phase Patterns of Neuronal Responses Reliably Discriminate Speech in Human Auditory Cortex. Neuron 54, 1001–1010 (2007).

42. Park, H., Ince, R. A. A., Schyns, P. G., Thut, G. & Gross, J. Frontal Top-Down Signals Increase Coupling of Auditory Low-Frequency Oscillations to Continuous Speech in Human Listeners. Curr. Biol. 25, 1649–1653 (2015).

43. Peelle, J. E., Gross, J. & Davis, M. H. Phase-locked responses to speech in human auditory cortex are enhanced during comprehension. Cereb. Cortex 23, 1378–1387 (2013).

44. Nourski, K. V. et al. Temporal Envelope of Time-Compressed Speech Represented in the Human Auditory Cortex. J. Neurosci. 29, 15564–15574 (2009).

45. Gross, J. et al. Speech Rhythms and Multiplexed Oscillatory Sensory Coding in the Human Brain. PLoS Biol. 11, (2013).

46. Ng, B. S. W., Schroeder, T. & Kayser, C. A Precluding But Not Ensuring Role of Entrained Low-Frequency Oscillations for Auditory Perception. J. Neurosci. 32, 12268–12276 (2012).

47. Ding, N. & Simon, J. Z. Cortical entrainment to continuous speech: functional roles and interpretations. Front. Hum. Neurosci. 8, (2014).

48. Makeig, S. Dynamic Brain Sources of Visual Evoked Responses. Science (80-. ). 295, 690–694 (2002).

49. Sauseng, P. et al. Are event-related potential components generated by phase resetting of brain oscillations? A critical discussion. Neuroscience 146, 1435–1444 (2007).

50. Liegeois-Chauvel, C., Musolino, A. & Chauvel, P. Localization of the primary auditory area in man. Brain 114A, 139–153 (1991).

51. Howard, M. A. et al. Auditory cortex on the human posterior superior temporal gyrus. J. Comp. Neurol. 416, 79–92 (2000).

52. Brugge, J. F. et al. Functional localization of auditory cortical fields of human: Click-train stimulation. Hear. Res. 238, 12–24 (2008).

53. Schroeder, C. E., Lakatos, P., Kajikawa, Y., Partan, S. & Puce, A. Neuronal oscillations and visual amplification of speech. Trends Cogn. Sci. 12, 106–113 (2008).

54. Capilla, A., Pazo-Alvarez, P., Darriba, A., Campo, P. & Gross, J. Steady-state visual evoked potentials can be explained by temporal superposition of transient event-related responses. PLoS One 6, (2011).

55. Keitel, C., Quigley, C. & Ruhnau, P. Stimulus-Driven Brain Oscillations in the Alpha Range: Entrainment of Intrinsic Rhythms or Frequency-Following Response? J. Neurosci. 34, 10137–10140 (2014).

56. VanRullen, R., Zoefel, B. & Ilhan, B. On the cyclic nature of perception in vision versus audition. Philos. Trans. R. Soc. B Biol. Sci. 369, (2014).

57. Zoefel, B. & Heil, P. Detection of near-threshold sounds is independent of eeg phase in common frequency bands. Front. Psychol. 4, (2013).

58. Howard, M. F. & Poeppel, D. Discrimination of Speech Stimuli Based on Neuronal Response Phase Patterns Depends on Acoustics But Not Comprehension. J. Neurophysiol. 104, 2500–2511 (2010).

59. Szymanski, F. D., Rabinowitz, N. C., Magri, C., Panzeri, S. & Schnupp, J. W. H. The Laminar and Temporal Structure of Stimulus Information in the Phase of Field Potentials of Auditory Cortex. J. Neurosci. 31, 15787–15801 (2011).

60. Henry, M. J. & Obleser, J. Frequency modulation entrains slow neural oscillations and optimizes human listening behavior. Proc. Natl. Acad. Sci. 109, 2009–100 (2012).

61. Wang, X.-J. Neurophysiological and Computational Principles of Cortical Rhythms in Cognition. Physiol. Rev. (2010). doi:10.1152/physrev.00035.2008

62. Cannon, J. et al. Neurosystems: Brain rhythms and cognitive processing. European Journal of Neuroscience (2014). doi:10.1111/ejn.12453

63. Walker, K. M. M., Bizley, J. K., King, A. J. & Schnupp, J. W. H. Multiplexed and Robust Representations of Sound Features in Auditory Cortex. J. Neurosci. (2011). doi:10.1523/JNEUROSCI.2074-11.2011

64. Fontolan, L., Morillon, B., Liegeois-Chauvel, C. & Giraud, A. L. The contribution of frequency-specific activity to hierarchical information processing in the human auditory cortex. Nat. Commun. (2014). doi:10.1038/ncomms5694

65. Kayser, S. J., Ince, R. A. A., Gross, J. & Kayser, C. Irregular Speech Rate Dissociates Auditory Cortical Entrainment, Evoked Responses, and Frontal Alpha. J. Neurosci. (2015). doi:10.1523/JNEUROSCI.2243-15.2015

66. Fries, P. A mechanism for cognitive dynamics: Neuronal communication through neuronal coherence. Trends in Cognitive Sciences (2005). doi:10.1016/j.tics.2005.08.011

67. Michalareas, G. et al. Alpha-Beta and Gamma Rhythms Subserve Feedback and Feedforward Influences among Human Visual Cortical Areas. Neuron 89, 384–397 (2016).

68. Buzsáki, G., Anastassiou, C. a. & Koch, C. The origin of extracellular fields and currents – EEG, ECoG, LFP and spikes. Nat. Rev. Neurosci. 13, 407–420 (2012).

69. Hopfield, J. J. Encoding for computation: Recognizing brief dynamical patterns by exploiting effects of weak rhythms on action-potential timing. Proc. Natl. Acad. Sci. (2004). doi:10.1073/pnas.0401125101

70. Arieli, A., Sterkin, A., Grinvald, A. & Aertsen, A. Dynamics of ongoing activity: Explanation of the large variability in evoked cortical responses. Science (80-. ). (1996). doi:10.1126/science.273.5283.1868

71. Super, H., van der Togt, C., Spekreijse, H. & Lamme, V. A. Internal state of monkey primary visual cortex (V1) predicts figure-ground perception. J Neurosci (2003).

72. Fiser, J., Chiu, C. & Weliky, M. Small modulation of ongoing cortical dynamics by sensory input during natural vision. Nature (2004). doi:10.1038/nature02907

73. Monto, S., Palva, S., Voipio, J. & Palva, J. M. Very Slow EEG Fluctuations Predict the Dynamics of Stimulus Detection and Oscillation Amplitudes in Humans. J. Neurosci. (2008). doi:10.1523/JNEUROSCI.1910-08.2008

74. Busch, N. A., Dubois, J. & VanRullen, R. The Phase of Ongoing EEG Oscillations Predicts Visual Perception. J. Neurosci. (2009). doi:10.1523/JNEUROSCI.0113-09.2009

75. Mathewson, K. E., Gratton, G., Fabiani, M., Beck, D. M. & Ro, T. To see or not to see: prestimulus alpha phase predicts visual awareness. J Neurosci (2009). doi:10.1523/JNEUROSCI.3963-08.2009

76. Neuling, T., Rach, S., Wagner, S., Wolters, C. H. & Herrmann, C. S. Good vibrations: Oscillatory phase shapes perception. Neuroimage (2012). doi:10.1016/j.neuroimage.2012.07.024

77. Jensen, O., Bonnefond, M. & VanRullen, R. An oscillatory mechanism for prioritizing salient unattended stimuli. Trends in Cognitive Sciences (2012). doi:10.1016/j.tics.2012.03.002

78. Nobre, A., Correa, A. & Coull, J. The hazards of time. Current Opinion in Neurobiology (2007). doi:10.1016/j.conb.2007.07.006

79. Andreou, L. V., Kashino, M. & Chait, M. The role of temporal regularity in auditory segregation. Hear. Res. (2011). doi:10.1016/j.heares.2011.06.001

80. Jones, M. R., Moynihan, H., Mackenzie, N. & Puente, J. Temporal Aspects of Stimulus-Driven Attending in Dynamic Arrays. Am. Psychol. Soc. 13, 313–319 (2002).

81. Hickok, G., Farahbod, H. & Saberi, K. The Rhythm of Perception: Entrainment to Acoustic Rhythms Induces Subsequent Perceptual Oscillation. Psychol. Sci. 26, 1006–1013 (2015).

82. Creutzfeldt, O., Ojemann, G. & Lettich, E. Neuronal activity in the human lateral temporal lobe II. Responses to the subjects own voice. Exp. brain Res. (1989). doi:10.1007/BF00249600

83. Houde, J. F. & Jordan, M. I. Sensorimotor adaptation of speech I: Compensation and adaptation. J. Speech, Lang. Hear. Res. (2002). doi:10.1044/1092-4388(2002/023)

84. Held, R. & Hein, A. V. Adaptation of disarranged hand-eye coordination contingent upon reafferent stimulation. Percept. Mot. Ski. (1958). doi:10.2466/pms.8.3.87-90

85. Müller-Preuss, P. & Ploog, D. Inhibition of auditory cortical neurons during phonation. Brain Res. (1981). doi:10.1016/0006-8993(81)90491-1

86. Flinker, A. et al. Single-Trial Speech Suppression of Auditory Cortex Activity in Humans. J. Neurosci. (2010). doi:10.1523/JNEUROSCI.1809-10.2010

87. Fukuda, M. et al. Cortical gamma-oscillations modulated by listening and overt repetition of phonemes. Neuroimage (2010). doi:10.1016/j.neuroimage.2009.10.047

88. Towle, V. L. et al. ECoG gamma activity during a language task: Differentiating expressive and receptive speech areas. Brain (2008). doi:10.1093/brain/awn147

89. Crone, N. E., Boatman, D., Gordon, B. & Hao, L. Induced electrocorticographic gamma activity during auditory perception. Brazier Award-winning article, 2001. Clin. Neurophysiol. (2001). doi:10.1016/S1388-2457(00)00545-9

90. Eliades, S. J. & Wang, X. Sensory-motor interaction in the primate auditory cortex during self-initiated vocalizations. J. Neurophysiol. 89, 2194–2207 (2003).

91. Wada, J. & Rasmussen, T. Intracarotid Injection of Sodium Amytal for the Lateralization of Cerebral Speech Dominance. J. Neurosurg. 106, 1117–1133 (2007).

92. Ellmore, T. M. et al. Temporal lobe white matter asymmetry and language laterality in epilepsy patients. Neuroimage 49, 2033–2044 (2010).

93. Conner, C. R., Ellmore, T. M., Pieters, T. a, Disano, M. a & Tandon, N. Variability of the relationship between electrophysiology and BOLD-fMRI across cortical regions in humans. J. Neurosci. 31, 12855–65 (2011).

94. Tandon, N. Mapping of Human Language. Clin. Brain Mapp. 203–218 (2012).

95. Oldfield, R. C. The assessment and analysis of handedness: The Edinburgh inventory. Neuropsychologia 9, 97–113 (1971).

96. Hamberger, M. J. & Seidel, W. T. Auditory and visual naming tests: normative and patient data for accuracy, response time, and tip-of-the-tongue. J. Int. Neuropsychol. Soc. 9, 479–489 (2003).

97. Ellmore, T. M., Beauchamp, M. S., O’Neill, T. J., Dreyer, S. & Tandon, N. Relationships between essential cortical language sites and subcortical pathways. J Neurosurg 111, 755–766 (2009).

98. Dale, A. M., Fischl, B. & Sereno, M. I. Cortical surface-based analysis. I. Segmentation and surface reconstruction. Neuroimage 9, 179–94 (1999).

99. Cox, R. W. AFNI: Software for analysis and visualization of functional magnetic resonance neuroimages. Comput Biomed Res 29, 162–173 (1996).

100. Pieters, T. A., Conner, C. R. & Tandon, N. Recursive grid partitioning on a cortical surface model: an optimized technique for the localization of implanted subdural electrodes. J. Neurosurg. 118, 1086–1097 (2013).

101. Gonzalez-Martinez, J. et al. Stereotactic placement of depth electrodes in medically intractable epilepsy. J. Neurosurg. 120, 639–644 (2014).

102. Gonzalez-Martinez, J. et al. Technique, Results, and Complications Related to Robot-Assisted Stereoelectroencephalography. Neurosurgery 78, 169–180 (2016).

103. Tandon, N. in Textbook of Epilepsy Surgery (ed. Luders, H.) 1001–1015 (McGraw Hill, 2008). at <http://www.crcnetbase.com/doi/abs/10.3109/9780203091708-129>

104. Bruns, a, Eckhorn, R., Jokeit, H. & Ebner, a. Amplitude envelope correlation detects coupling among incoherent brain signals. Neuroreport 11, 1509–1514 (2000).

105. Kadipasaoglu, C. M. et al. Development of grouped icEEG for the study of cognitive processing. Front. Psychol. 6, 1–7 (2015).

106. Whaley, M. L., Kadipasaoglu, C. M., Cox, S. J. & Tandon, N. Modulation of Orthographic Decoding by Frontal Cortex. J. Neurosci. 36, 1173–1184 (2016).

107. Conner, C. R., Chen, G., Pieters, T. A. & Tandon, N. Category specific spatial dissociations of parallel processes underlying visual naming. Cereb. Cortex 24, 2741–2750 (2014).

108. Berry, M. W., Browne, M., Langville, A. N., Pauca, V. P. & Plemmons, R. J. Algorithms and applications for approximate nonnegative matrix factorization. Comput. Stat. Data Anal. (2007). doi:10.1016/j.csda.2006.11.006

109. Argall, B. D., Saad, Z. S. & Beauchamp, M. S. Simplified intersubject averaging on the cortical surface using SUMA. Hum. Brain Mapp. 27, 14–27 (2006).

110. Fischl, B., Sereno, M. I., Tootell, R. B. H. & Dale, a M. High-resolution inter-subject averaging and a surface-based coordinate system. Hum. Brain Mapp. 8, 272–284 (1999).

111. Saad, Z. S. & Reynolds, R. C. Suma. Neuroimage 62, 768–773 (2012).

112. Miller, K. J. et al. Spectral Changes in Cortical Surface Potentials during Motor Movement. J. Neurosci. 27, 2424–2432 (2007).

113. Esposito, F. et al. Cortex-based inter-subject analysis of iEEG and fMRI data sets: Application to sustained task-related BOLD and gamma responses. Neuroimage 66, 457–468 (2013).

114. Kadipasaoglu, C. M. et al. Surface-based mixed effects multilevel analysis of grouped human electrocorticography. Neuroimage 101, 215–224 (2014).

